# Phagocytosis of primary human macrophages is elevated by ex vivo supplementation with n-3 PUFA

**DOI:** 10.1101/2025.07.30.667640

**Authors:** Rebecca Kirchhoff, Michel André Chromik, Nils Helge Schebb

## Abstract

**Scope:** Epidemiologic studies show that a high n-3 polyunsaturated fatty acid (PUFA) status is beneficial for health and inflammatory diseases. However, results of nutrition studies investigating the impact of n-3 PUFA intake on immune functions, such as phagocytosis, are contradictory. In order to gain more insights into the role of n-3 PUFAs on phagocytosis, we investigated the modulation of phagocytosis by n-3 PUFAs and derived oxylipins in human macrophages.

**Methods and results:** Using an established ex vivo supplementation strategy, primary human macrophages were supplemented with docosahexaenoic acid (DHA) and eicosapentaenoic acid (EPA). The PUFA pattern of the cells was shifted from a low n-3 PUFA status towards a high n-3 PUFA status. This was accompanied by a shift in the oxylipin pattern, reduced pro-inflammatory prostaglandin levels, increased phagocytosis in the supplemented macrophages, and reduced inhibitory effect of PGE_2_ on phagocytosis. However, when tested alone, n-3 PUFA derived oxylipins did not impact phagocytosis.

**Conclusion:** Under controlled conditions, an increased n-3 PUFA status of macrophages resulted in an elevation of phagocytosis. Less formation of prostaglandins could contribute to this effect, whereas n-3 PUFA derived oxylipins, particularly multihydroxy PUFAs, appear to have a limited impact on phagocytosis following n-3 PUFA supplementation.

## Introduction

Dietary intake of polyunsaturated fatty acids (PUFAs) is vital for humans and a high n-3 PUFA status is associated with beneficial health outcomes.^[1]^ Whereas the typical diet in Europe and the US includes large amounts of n-6 PUFAs, the intake of n-3 PUFAs – especially long-chain n-3 PUFAs eicosapentaenoic acid (EPA) and docosahexaenoic acid (DHA) – is low.^[2, 3]^ In contrast, a high n-3 PUFA status, e.g. by supplementing n-3 PUFAs from fish or algae oil, results in a lower %n-6 in HUFA, a biomarker associated with cardiovascular diseases.^[4]^ However, the underlying mechanisms of action by which n-3 PUFAs affect biological functions remain to be elucidated.

PUFAs are important components of the cell membranes affecting fluidity, membrane-associated proteins, formation of lipid rafts and the pattern of formed eicosanoids and other oxylipins.^[5]^ Several oxylipins are lipid mediators involved in the regulation of physiological and inflammatory processes: n-6 PUFA derived oxylipins such as prostaglandins (PG) formed by cyclooxygenase (COX) activity and leukotrienes formed by 5-lipoxygenase (LOX) activity from arachidonic acid (ARA) are well-investigated pro-inflammatory mediators^[6, 7]^ causing e.g., pain, fever and asthma.^[8]^ Increased intake of n-3 PUFAs shifts the oxylipin pattern by a reduced conversion of n-6 PUFAs, and an increased in the formation of n-3 PUFA derived oxylipins.^[9]^ Consequently, oxylipins are likely to contribute to the beneficial effects of n-3 PUFAs on immune functions such as phagocytosis.

Phagocytosis is a key mechanism of innate host defense directly neutralizing pathogens. Additionally, phagocytosis plays a role in the active resolution of inflammation by removing apoptotic cells and cell debris, thereby enabling restoration of healthy tissue.^[10]^ Several oxylipins, especially multihydroxy oxylipins derived from EPA and DHA, are discussed to have anti-inflammatory effects, such as modulation of phagocytosis,^[11, 12]^ but their formation and presence in human macrophages remains controversial.^[13]^

In order to elucidate how n-3 PUFAs impact phagocytosis and whether n-3 PUFA derived oxylipins contribute to their effects, primary human ‘inflammatory’ M1- and ‘anti-inflammatory’ M2-like macrophages, representing extremes of macrophage phenotypes, were supplemented with DHA and EPA using an established ex vivo strategy.^[14]^ Changes in FA and oxylipin pattern of macrophages following n-3 PUFA supplementation were quantified by means of LC-MS/MS. Phagocytic activity following n-3 PUFA supplementation as well as the impact of n-3 PUFA derived oxylipins, which were increased by the supplementation, were investigated in both phenotypes of primary macrophages.

## Experimental section

### Chemicals and biological material

Non-fasted human AB plasma was obtained from the blood donation center at University Hospital Düsseldorf (Düsseldorf, Germany). Lymphocyte separation medium 1077 was purchased from PromoCell (Heidelberg, Germany). Recombinant human GM-CSF, M-CSF, IFNγ and IL-4 produced in *Escherichia coli* were bought from Thermo Fisher Scientific (Langenselbold, Germany). RPMI 1640 cell culture medium, L-glutamine and penicillin/streptomycin (5.000 units penicillin and 5 mg/mL streptomycin), LPS from *E. coli* (0111:B4), dextran 500 from *Leuconostoc spp.* and copper sulfate pentahydrate were from Merck (Taufkirchen, Germany). Docosahexaenoic acid (DHA > 99%) and eicosapentaenoic acid (EPA > 99%) were bought from Nu-check Prep, Inc. (Elysian, Minnesota, United States). Prostaglandin E_2_ (PGE_2_), 5-HEPE, 4-HDHA, 7-HDHA, 12(*S*)-HEPE, 15(*S*)-HEPE, 14(*S*)-HDHA, 17(*S*)-HDHA, 17(18)-EpETE and 19(20)-EpDPE were bought from Cayman Chemical (local distributor Biomol, Hamburg, Germany). Accutase, DMSO and formaldehyde were purchased at Carl Roth (Karlsruhe, Germany). Fluorescence-labeled polystyrene microspheres (latex beads) were obtained as 2% solution from Thermo Fisher Scientific (Langenselbold, Germany). BCA reagent A was bought from Fisher Scientific (Schwerte, Germany). The ultra-pure water with a conductivity of >18 MΩ*cm was generated by the Barnstead Genpure Pro system from Thermo Fisher Scientific (Langenselbold, Germany).

### Isolation and differentiation of primary human macrophages

Primary human macrophages were isolated as described.^[15]^ In brief, peripheral blood mononuclear cells (PBMCs) were isolated from buffy coats provided by blood donations at the University Hospital Düsseldorf, Germany or at Deutsches Rotes Kreuz West, Hagen, Germany. The blood donations were drawn with the informed consent of healthy human subjects and the study was approved by the Ethical Committee of the University of Wuppertal. PBMC were isolated by dextran (5%) sedimentation for 30 min followed by centrifugation (800 × *g* without deceleration, 10 min, 20 °C) on lymphocyte separation medium. The leucocyte ring was isolated and washed twice with PBS. The cells were seeded in 60.1 mm² dishes in RPMI medium (100 IU/mL penicillin and 100 µg/mL streptomycin (P/S), 2 mM L-glutamine) in a humidified incubator at 37 °C and 5% CO_2_ for 1-2 h. Non-adherent cells were removed by gently washing, and RPMI medium (P/S, 2 mM L-glutamine) supplemented with 5% human AB plasma was added to the cells. Using an established protocol,^[15]^ cells were polarized towards M1-like macrophages using 10 ng/mL granulocyte-macrophage colony-stimulating factor (GM-CSF) for 7 days and additional 10 ng/mL interferon γ (IFNγ) for the final 48 h or towards M2-like macrophages using 10 ng/mL macrophage (M)-CSF for 7 days and additional 10 ng/mL interleukin 4 (IL-4) for the final 48 h.

### Supplementation of human macrophages with n-3 PUFA

Supplementation experiments were carried out as described.^[14]^ In brief, for the selection of a suitable plasma for the cell culture medium, five non-fasting human AB plasmas were obtained at the University Hospital Düsseldorf, Germany and analyzed for fatty acid levels and oxylipin concentrations **(Figure S1, Tables S1, S2)**. Plasmas C and E were pooled (pool 1) resulting in a plasma which fulfilled all criteria:^[14]^ %n-6 in HUFA >75%, n-3 PUFA <0.25 mM, total FA <10 mM, low oxylipin concentrations.

For selection of suitable subjects/buffy coats which reflect the poor n-3 PUFA status of average subjects in Europe and the US, only subjects having a %n-6 in HUFA ≥70% in erythrocytes were selected for the supplementation experiments **(Table S3)**.^[14]^

For supplementation, following predilution in DMSO (50 mM), EPA and DHA were slowly mixed with human plasma and added (5% (*v*/*v*)) to RPMI medium (P/S, 2 mM L-glutamine) (10.3 µM DHA, 5.5 µM EPA) **(Figure 1)**. Medium containing the same plasma, but without added DHA and EPA, served as control medium with low n-3 PUFA content (low). For n-3 PUFA supplementation, macrophages were incubated with supplemented medium (high) for 2 days. Macrophages were harvested by cold shock method,^[15]^ and cell pellets were frozen at - 80 °C until analysis.

**Figure 1:**
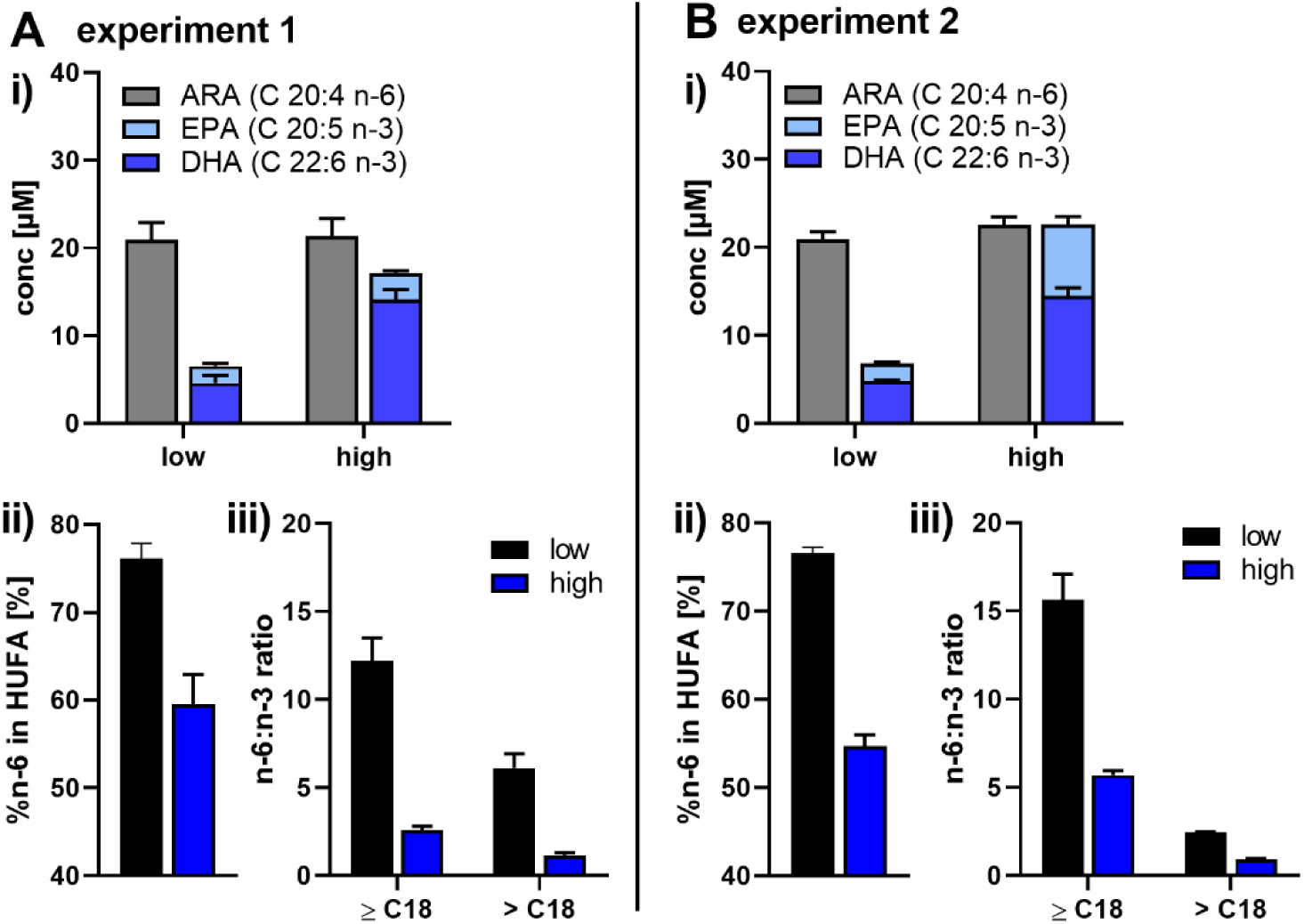
The FA pattern of cell culture medium is changed by addition of n-3 PUFAs. In two independent experiments **(A, B)**, medium was prepared by using 5% (*v*/*v*) plasma pool 1 (low). For the supplemented medium (high), 10.3 µM DHA (>99%) and 5.5 µM EPA (>99%) were added. **i)** Total FA concentrations were determined by means of LC-MS/MS. **ii)** %n-6 in HUFA and **iii)** n-6:n-3 ratios were determined based on total FA concentrations. Results are shown as mean ± SD, *n* = 3.

### Phagocytosis assay

Phagocytosis was carried out as described.^[16]^ Briefly, differentiated and supplemented macrophages were harvested using accutase and transferred into 96 well plates at 50,000 cells/well. After resting overnight, cells were preincubated with test substances or vehicle control as indicated. For phagocytosis, supernatants were discarded and fluorescence-labeled polystyrene microspheres (beads) diluted in medium (6.25 ·10^-3^% beads, 5% human plasma) were added together with the preincubated test substance to the cells for 2 h. Phagocytosis was stopped by discarding the supernatants, carefully washing the cells with PBS and fixating the cells with 4% formaldehyde. Fluorescence of beads was measured using a plate reader (Infinite 200 PRO, Tecan) (λ_ex_ 535 nm, λ_em_ 575 nm). For DAPI staining 10 µg/mL DAPI was added to the cells for 15 min at room temperature followed by three times washing with PBS and fluorescence measurement using the plate reader (λ_ex_ 358 nm, λ_em_ 461 nm).

### Quantification of fatty acyl and oxylipin levels by means of LC-MS/MS

Oxylipin determination was carried out in cell pellets, medium and human plasma as described.^[17–20]^ Briefly, cell pellets were resuspended in MeOH/H_2_O (50/50, *v*/*v*), and sonicated. After addition of internal standards (IS), proteins were precipitated using MeOH (for non-esterified oxylipins) or *iso*-propanol (for total oxylipins) followed by centrifugation (20 000 × *g*, 10 min, 4 °C). For quantification of total oxylipin levels, supernatants were saponified using 0.6 M KOH in MeOH/H_2_O (75/25, *v*/*v*) for 30 min at 60 °C. Oxylipins were purified using solid phase extraction (SPE) and analyzed by means of LC–MS/MS using an 1290 Infinity II LC system (Agilent), coupled to a QTRAP mass spectrometer operated in electrospray ionization (ESI(-)) mode (Sciex, Darmstadt, Germany) operated in scheduled selected reaction monitoring. Fatty acyl concentrations from the same samples as for the total oxylipins were determined by LC-MS/MS as described.^[21]^ Oxylipin and fatty acyl concentrations were quantified using external calibrations with IS.

Analyst (Sciex, version 1.7) was used for instrument control and Multiquant software (Sciex, version 3.0.2) was used for data analysis. The concentrations of oxylipins and fatty acids in the macrophages were calculated based on the protein content determined via bicinchoninic acid assay.^[22]^

### Data analysis

Data are presented as mean ± standard deviation (SD). Statistical analysis and visualization of data was performed using the Prism software (GraphPad Software, version 8.4., San Diego, CA, USA). For statistical analysis, 1-way ANOVA following Dunnett’s or Sidak’s multiple comparison tests were performed as indicated in the figure captions. Statistical significance is depicted as * *p* ≤0.05, ** *p* ≤0.01, *** *p* ≤0.001. %n-6 in HUFA and n-6:n-3 ratios were calculated from total FA concentrations:^[23]^

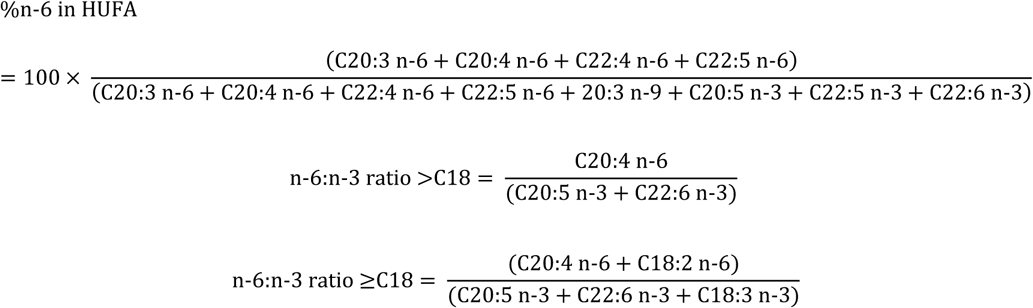

## Results

In order to better understand the effects of long-chain n-3 PUFAs and their oxylipins on human immune cells, a reliable ex vivo supplementation strategy was used allowing to alter the FA and oxylipin patterns of primary human macrophages comparable to human intervention studies.^[14]^ Using an established assay for phagocytosis,^[16]^ changes in phagocytic activity following n-3 PUFA supplementation were investigated in M1- and M2-like macrophages.

### Ex vivo supplementation of primary human macrophages with n-3 PUFAs

The n-3 PUFA supplementation was carried out in two independent experiments, each with four human subjects. In order to enable a reproducible supplementation, key parameters for selection of buffy coats, plasma, n-3 PUFAs and composition of cell culture media^[14]^ were controlled **(Figures S1, S2 S3; Tables S1, S2, S3)**. The cell culture medium was prepared using 5% (*v*/*v*) of plasma (non-supplemented control, n-3 PUFA low) and plasma with 10.3 µM DHA and 5.5 µM EPA (n-3 PUFA high). This resulted in equal concentrations of ARA and EPA + DHA in the medium, a decrease of the %n-6 in HUFA by approximately 20%, and of the n-6:n-3 ratio by factor 10 **(Figure 1)**. For supplementation of the cells, macrophages were incubated with the n-3 PUFA supplemented medium (high) for two days.

n-3 PUFA supplementation decreased %n-6 in HUFA of macrophages from 85 ± 2% to 61 ± 4% (experiment 1) and 85 ± 1% to 56 ± 1% (experiment 2) **(Figure 2 (i))**. The relative amount of n-3 PUFAs, especially DHA and EPA, was increased in the cells by approximately 4-7%, whereas n-6 PUFAs such as ARA were decreased by approximately 3%; MUFA and SFA were not or only hardly changed **(Figure 2 (ii-iii))**. Similar to the changes in the FA pattern, oxylipins derived from EPA and DHA increased (e.g., 4-HDHA by approx. 2-3fold; 5-HETE by approx. 2-6fold) whereas ARA-derived oxylipins were decreased (e.g., 5-HETE by approx. 25%) in a comparable manner in both macrophage types **(Figure 3)**. However, 15-LOX products were more affected in the M2-like phenotype: e.g., 17-HDHA was increased by approx. 3-10fold by supplementation in M2-like, but not changed in M1-like; 15-HETE was decreased by 60-70% in M2-like, but only 20-30% in M1-like. Dihydroxy-PUFAs such as 5,15-diHETE or 5,15-diHEPE were below LLOQ. A clear decrease of COX products from ARA such as the PGE_2_ breakdown product 12-HHTrE (by approx. 60-70%) and prostaglandins (e.g., PGE_2_ by approx. 55-70% in experiment 1) was detected in both macrophage phenotypes following supplementation with n-3 PUFA **(Figure 3)**.

**Figure 2:**
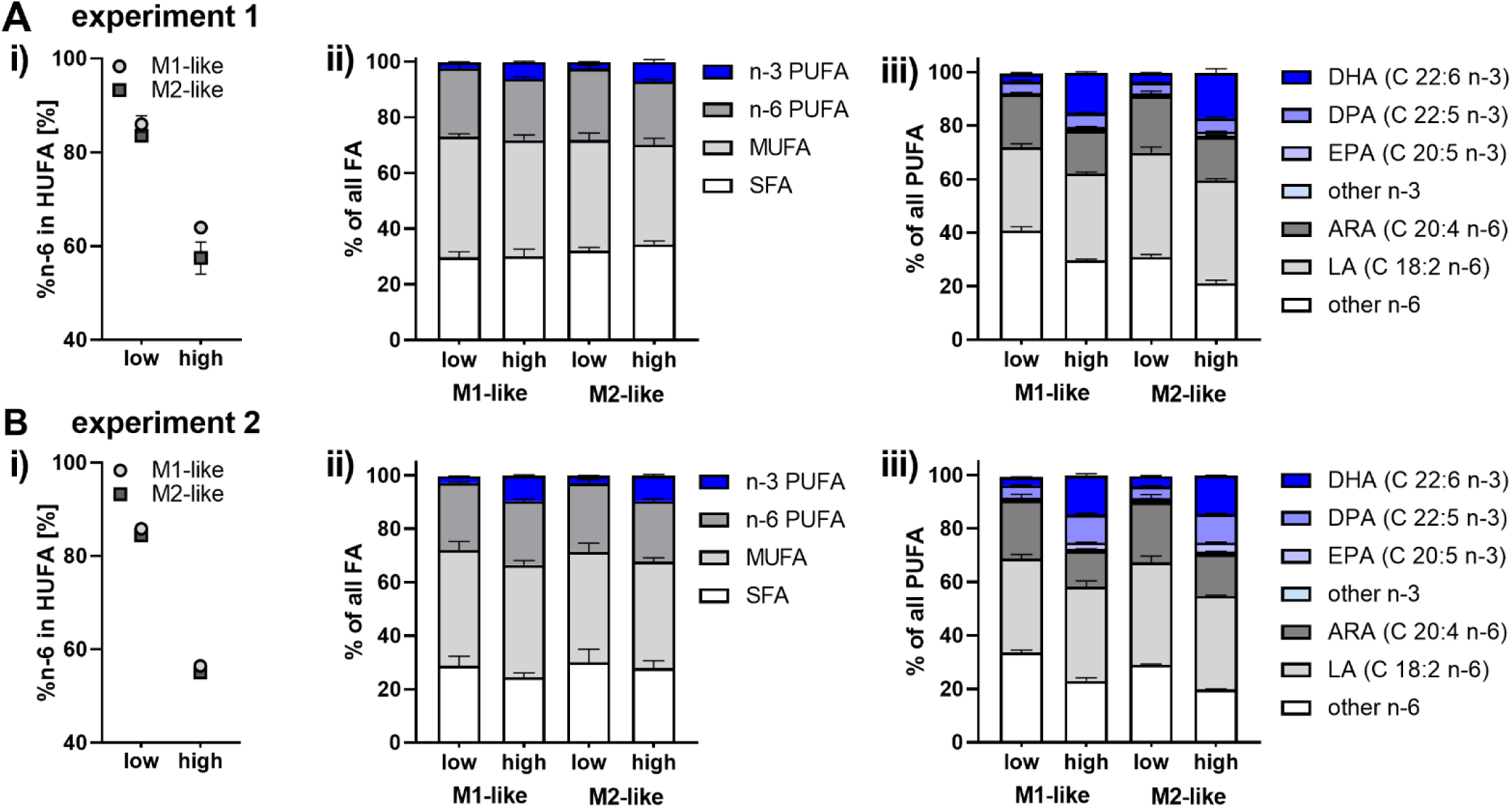
FA pattern in macrophages is shifted towards cells with a high n-3 PUFA status following supplementation with n-3 PUFA in both supplementation experiments (A, B). After isolation of monocytes, cells were differentiated into macrophages and supplemented using medium with 5% (*v*/*v*) plasma pool 1 (low) or medium containing plasma pool 1, 10.3 µM DHA (>99%) and 5.5 µM EPA (>99%) (high) for 2 days. Total FA were quantified in the cell pellets by means of LC-MS/MS. **i)** %n-6 in HUFA of macrophages. **ii)** Relative distribution of n-3 PUFA, n-6 PUFA, MUFA and SFA. **iii)** Relative distribution of PUFAs. Results are shown as mean ± SD, *n* = 3 pooled cells of 4 subjects for each experiment.

**Figure 3:**
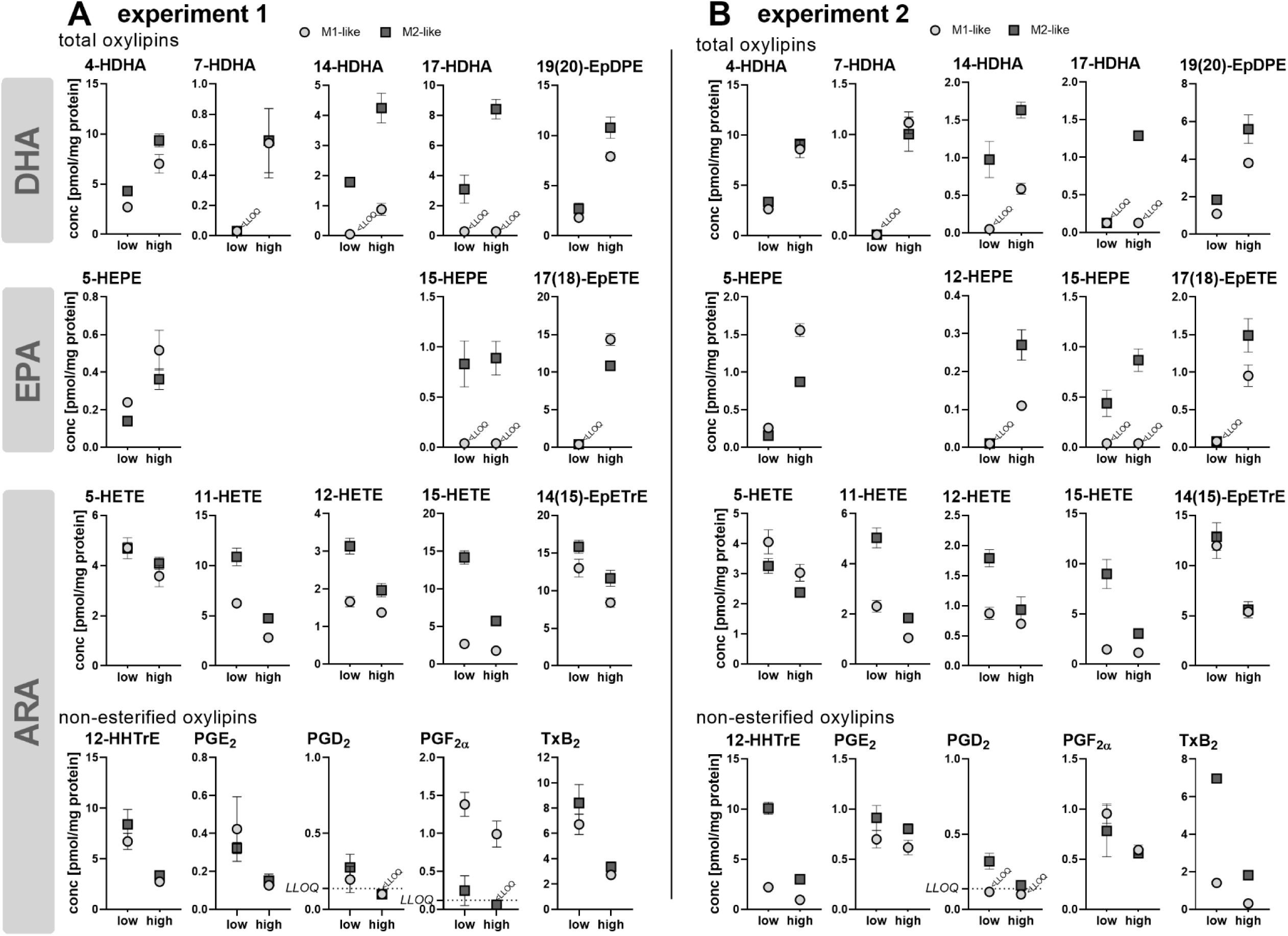
DHA and EPA derived oxylipins are increased, ARA derived oxylipins decreased in macrophages after supplementation with n-3 PUFA. In two independent experiments **(A, B)**, differentiated macrophages were supplemented using medium with 5% (*v*/*v*) plasma pool 1 (low) or medium containing plasma pool 1, 10.3 µM DHA (>99%) and 5.5 µM EPA (>99%) (high) for 2 days. Total oxylipins (monohydroxy-PUFA, epoxy-PUFA) and non-esterified oxylipins (12-HHTrE, prostaglandins, TxB_2_) were quantified in the cell pellets by means of LC-MS/MS. Concentrations of selected, most abundant oxylipins derived from DHA, EPA and ARA are shown as mean ± SD, *n* = 3 pooled cells of 4 subjects for each experiment.

### Ex-vivo supplementation with n-3 PUFA increases phagocytosis in macrophages

Phagocytic activity of non-supplemented (low) and n-3 PUFA supplemented (high) macrophages was compared using an established phagocytosis assay with fluorescent labeled polystyrene beads.^[16]^ Phagocytosis increased slightly but significantly in both macrophage phenotypes following n-3 PUFA supplementation **(Figure 4A, Figure S3)**. The increase in phagocytosis was more pronounced for the M1-like phenotype (e.g., for experiment 1: M1-like 117 ± 9%, M2-like 108 ± 12%).

**Figure 4:**
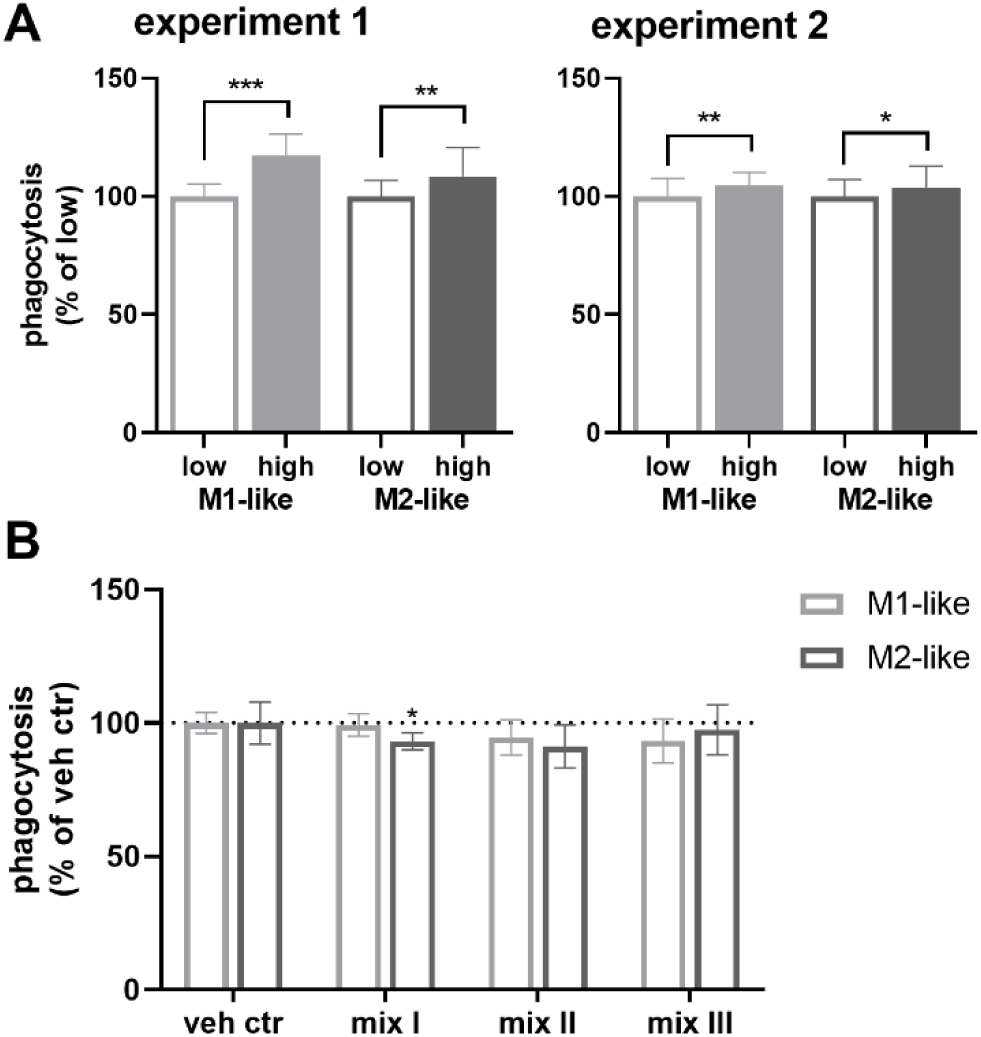
n-3 PUFA supplementation, but not the n-3 PUFA derived oxylipins increase phagocytosis in macrophages. **A)** After isolation of monocytes, cells were differentiated into macrophages and supplemented using medium with 5% (*v*/*v*) plasma pool 1 (low) medium containing plasma pool 1, 10.3 µM DHA (>99%) and 5.5 µM EPA (>99%) (high) for 2 days. Shown is phagocytosis as % change from the non-supplemented cells (low). Summarized results of 4 independently performed experiments (Figure S3) are shown as mean ± SD for *n* = 26 (experiment 1) or *n* = 58-60 replicates (experiment 2) from a pool of 4 subjects for each supplementation experiment. **B)** Non-supplemented macrophages were incubated with 300 nM oxylipins (mix I: 5-HEPE, 4-HDHA, 7-HDHA; mix II: 12(*S*)-, 15(*S*)-HEPE, 14(*S*)-, 17(*S*)-HDHA; mix III: 17(18)-EpETE, 19(20)-EpDPE, Table S4) or 0.1% DMSO (vehicle control) for 1 h. Summarized results of two independently performed experiments for phagocytosis are shown as % change from vehicle control as mean ± SD for *n* = 9-10 replicates from a pool of 4 subjects (see Figure S4 for all results). Statistical analysis was performed by 1-way ANOVA followed by A) Sidak’s or B) Dunnett’s multiple comparison test. Differences from vehicle control were considered significant at *p* values ≤0.5 (*), ≤0.01 (**) or ≤0.001 (***).

### Impact of oxylipins on phagocytosis

In order to investigate if oxylipins contribute to the increase in phagocytosis following n-3 PUFA supplementation, those oxylipins **(Table S4)** were tested for their effect on phagocytosis which were increased by n-3 PUFA supplementation. However, the oxylipins had no or slightly inhibitory effects on phagocytosis in non-supplemented macrophages **(Figure 4B, Figure S4)**. After phagocytosis, concentrations of the oxylipins in the supernatants were quantified demonstrating that target concentrations were achieved throughout the assay **(Table S5)**.

PGE_2_ which is known to inhibit phagocytosis in human macrophages^[16]^ was found to be decreased in the supernatants of n-3 PUFA supplemented macrophages, both at baseline and after an inflammatory stimulus (0.1µg/mL LPS) **(Figure 5A, B)**. Interestingly, when PGE_2_ is added exogenously to the cells, phagocytosis is 30-70% less inhibited in the macrophages following n-3 PUFA supplementation **(Figure 5C, D)**. In order to test, if the n-3 PUFA derived oxylipins abrogate the inhibitory effect of PGE_2_ on phagocytosis, non-supplemented macrophages were incubated with PGE_2_ and different oxylipins **(Table S4)**; concentrations of oxylipins were checked in the supernatants after phagocytosis **(Table S6)**. However, the tested oxylipins did not abrogate the inhibitory effect of PGE_2_ on phagocytosis **(Figure 5E, Figure S5)**.

**Figure 5:**
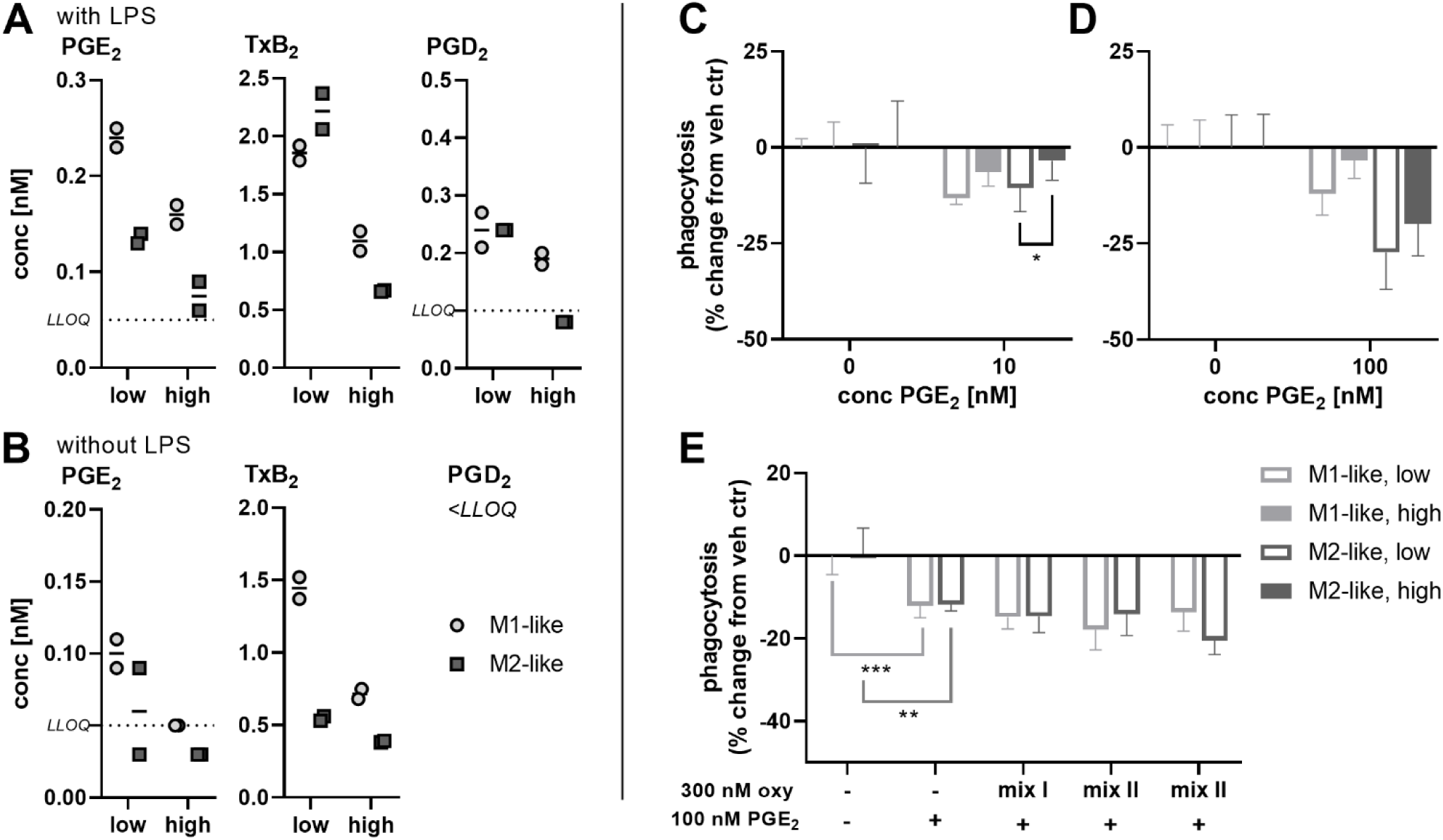
n-3 PUFA supplementation reduces inhibitory effect of PGE_2_ on phagocytosis. Macrophages were supplemented using medium with 5% (*v*/*v*) plasma pool 1 (low) or medium containing plasma pool 1, 10.3 µM DHA (>99%) and 5.5 µM EPA (>99%) (high) for 2 days. Cells were transferred into 96 well plates (50,000 cells per well) and incubated **A)** with 0.1 µg/mL LPS or **B)** without LPS for 4 h. Non-esterified oxylipin concentrations were quantified in the supernatants by means of LC-MS/MS. Concentrations of COX products are shown for one representative experiment. **(C-E)** For phagocytosis, cells were preincubated with **C)** 10 nM or **D)** 100 nM PGE_2_ for 1 h. Results of one representative experiment are shown as mean ± SD for *n* = 4-5 replicates from a pool of 4 subjects. **E)** Non-supplemented cells were preincubated with 100 nM PGE_2_ and 300 nM oxylipins (mix I: 5-HEPE, 4-HDHA, 7-HDHA; mix II: 12(*S*)-, 15(*S*)-HEPE, 14(*S*)-, 17(*S*)-HDHA; mix III: 17(18)-EpETE, 19(20)-EpDPE, Table S4) or 0.1% DMSO (vehicle control) for 1 h. Shown is phagocytosis as % change from vehicle control as mean ± SD for *n* = 3-10 replicates from a pool of 4 subjects. Statistical analysis was performed by 1-way ANOVA followed by Sidak’s multiple comparison test. Differences from vehicle control were considered significant at *p* values ≤0.5 (*), ≤0.01 (**) or ≤0.001 (***).

## Discussion

A sufficient intake of long-chain n-3 PUFAs such as EPA and DHA is required for human health, and associated with beneficial health effects.^[24, 25]^ Here, we investigated the impact of these PUFAs and their derived oxylipins on phagocytosis in human macrophages.

In order to enable a reliable investigation of the physiological effects of n-3 PUFAs without carrying out intervention studies, an ex vivo supplementation strategy was used,^[14]^ and key parameters for all steps of the supplementation were controlled: Briefly, (i) only subjects reflecting the poor n-3 PUFA status of the population of Europe and the US were included **(Table S3)**, (ii) a suitable plasma low in oxylipin concentrations, low total FA and n-3 PUFA concentrations – reflecting the low n-3 PUFA status of the average population in Europe and the US – was selected **(Figure S1, Tables S1, S2)**, (iii) only pure n-3 PUFAs were used (max. 0.5% (*w*/*w*) oxylipins), and (iv) cell culture media were always freshly prepared and replaced every 2-3 days in order to keep autoxidative formation of oxylipins low **(Figure S3)**.

Incubation of the macrophages with the n-3 PUFA enriched medium (high) **(Figure 1)** for 2 days remarkedly changed the FA profile **(Figure 2)** as well as the oxylipin pattern **(Figure 3)** of the macrophages: %n-6 in HUFA of non-supplemented macrophages (85 ± 2%) is comparable to values found in tissues from subjects having a low n-3 PUFA status,^[4]^ and was decreased to values between 55-64% by supplementation. Similar values for the %n-6 in HUFA were achieved in human nutrition in erythrocytes using oral supplementation and daily doses of 1-2 g.^[26–28]^ Whereas MUFA and SFA levels remained unchanged, n-6 PUFAs were replaced by n-3 PUFAs such as EPA and DHA **(Figure 2)** which is in accordance with results of supplementation studies in humans and animals.^[26–29]^

Similarly, the increase in n-3 PUFA derived oxylipins and decrease of ARA derived ones **(Figure 3)** is comparable to previous supplementation experiments with primary human M2-like macrophages,^[14]^ and consistent with nutrition intervention studies.^[29–32]^ For M1-like macrophages smaller changes in concentrations of 15-LOX products following supplementation were observed compared to the M2-like phenotype. This can be explained by the fact that this enzyme is not abundant in the M1-like phenotype, but strongly expressed in the M2-like phenotype.^[20, 33]^ Di-and multihydroxy PUFAs were not detected in the macrophages. This differs from previous studies,^[14, 34]^ and could be explained by a sample preparation using 50% (*v*/*v*) MeOH for dissolving the cell pellet which inhibits enzyme activity and prevents artificial formation of 15-LOX products during sample preparation.

Overall, the FA pattern of monocytes shifted from one comparable to subjects with a typical Western diet to one comparable to subjects with a high n-3 PUFA status. The results of both supplementation experiments are highly comparable which demonstrates the reproducibility of the ex vivo supplementation strategy.

In both independent supplementation experiments phagocytosis was increased in the n-3 PUFA supplemented macrophages **(Figure 4A, Figure S3)**. This is consistent with cell culture experiments showing that membrane FA composition is associated with altered phagocytic activity.^[35–37]^ However, nutrition studies in humans led to conflicting results: Whereas some studies reported an increase of phagocytosis by n-3 PUFA supplementation,^[38–40]^ most studies found no effect.^[41–45]^ Of note, results are difficult to compare as doses, duration and PUFAs used for supplementation as well as the method of measuring phagocytosis are highly different between the studies. The finding that n-3 PUFAs have no effect in nutrition studies could be related to the fact that studies included all kinds of subjects, with low and high n-3 PUFA status. However, consistent with their role as essential food ingredient, there is growing evidence that only subjects with a low n-3 PUFA status benefit from a supplementation with n-3 PUFA.^[46–48]^ Our results support this finding as only subjects are included with a low n-3 PUFA status, and PUFA profile and immune functions of macrophages are altered by supplementation.

The mechanisms of action by which n-3 PUFA influence phagocytosis are not fully understood: In addition to changes in membrane fluidity^[49]^ or altered gene expression,^[50]^ it is hypothesized that the shift of the lipid mediator pattern following n-3 PUFA supplementation contributes to the increase in phagocytosis.^[51]^ In order to investigate the impact of lipid mediators, n-3 PUFA derived oxylipins which were increased following n-3 PUFA supplementation **(Figure 3)** were tested for their impact on phagocytosis **(Table S4)**: 5-HEPE, 4- and 7-HDHA formed by 5-LOX activity and/or autoxidation^[52, 53]^ (mix I), 12-, 15-HEPE and 14-, 17-HDHA by 15-LOX activity^[54–56]^ (and autoxidation) (mix II) and the epoxy PUFA 17(18)-EpETE and 19(20)-EpDPE (mix III), which are formed by cytochrome P450 monooxygenases,^[57]^ but the increase here is caused by impurities in the PUFAs used for supplementation.^[14]^ Of note, the latter are partly converted into the corresponding *vic* dihydroxy PUFAs **(Table S5)**, presumably by conversion by soluble epoxide hydrolase activity. Using a relatively high concentration (300 nM) of the oxylipins did not impact phagocytosis **(Figure 4B, Figure S4)** suggesting that these n-3 PUFA derived oxylipins (alone) are not the reason for the increased phagocytosis following n-3 PUFA supplementation.

n-3 PUFA supplementation resulted in decreased levels of COX products such as prostaglandins in the macrophages **(Figure 3)** and supernatants **(Figure 5A, B)**. A reduced formation of prostaglandins, especially PGE_2_, could contribute to the increase of phagocytosis by supplementation, as PGE_2_ inhibits phagocytosis.^[16]^ Moreover, when PGE_2_ is added to the cells, phagocytosis is less inhibited in the n-3 PUFA supplemented cells **(Figure 5C, D)**. This could indicate that PGE_2_ acts differently on phagocytosis when combined with n-3 PUFA derived oxylipins. However, when testing n-3 PUFA derived oxylipins which were increased by n-3 PUFA supplementation **(Table S4)**, the oxylipins had no impact on the inhibitory effect of PGE_2_ on phagocytosis **(Figure 5E, Figure S5, Table S6)**. Recent studies have shown that 7(*S*),17(*S*)-DiHDHA (RvD5) affects the EP4 receptor through allosteric modulation, thereby reversing the inhibition of phagocytosis by PGE_2_ in murine macrophages.^[58]^ This fits to our results that PGE_2_ inhibits phagocytosis less in the n-3 PUFA supplemented cells. However, 7(*S*),17(*S*)-DiHDHA, which is discussed to modulate the EP4 receptor, was not detected in the macrophages. Similarly, other multihydroxy PUFA such as RvE2 and PD1, which were shown to increase phagocytosis in human macrophages,^[11, 12]^ could not be detected in the cells. This is consistent with several reports indicating that these compounds do not occur in macrophages.^[14, 34, 59]^ Moreover, formation and signaling of these multihydroxy PUFA in human macrophages remains controversial,^[13]^ indicating that these play only a minor role in increasing the phagocytosis following a n-3 PUFA supplementation. However, an increased phagocytosis is beneficial allowing an efficient defense against pathogens and the rapid clearance of apoptotic cells, both key processes contributing to tissue repair and homeostasis.^[10,60]^

Our findings support, that subjects with a low n-3 PUFA status – which is typical for people in Europe and the US when having a Western diet – benefit from a n-3 PUFA supplementation altering the PUFA profile and phagocytosis in the macrophages. Additionally, levels of pro-inflammatory prostaglandins are reduced which contributes to the beneficial effect of n-3 PUFAs. However, further research is needed elucidating the modes of action by which n-3 PUFAs and derived oxylipins impact phagocytosis and immune functions in humans.

## Conclusion

Increasing the n-3 PUFA status of primary human macrophages elevates phagocytosis, indicating an improved potential for the clearance of pathogens and cell debris. This beneficial effect of an n-3 PUFA supplementation could partly be explained by the reduced formation of pro-inflammatory prostaglandins such as PGE_2_. The n-3 PUFA derived oxylipins, increased by n-3 PUFA supplementation, alone had no effect. Further studies are needed to understand the mechanisms by which n-3 PUFAs elevate phagocytosis, involving the investigation of membrane fluidity, lipid rafts and membrane-associated proteins.

## Acknowledgments

We thank Katinka Langenohl for excellent technical support.

## Author contributions

**Rebecca Kirchhoff**: Conceptualization, Formal analysis, Investigation, Methodology, Writing – original draft, Writing – review and editing. **Michel André Chromik**: Formal analysis, Investigation, Writing – review and editing. **Nils Helge Schebb**: Conceptualization, Resources, Methodology, Funding acquisition, Writing – original draft, Writing – review and editing.

## Ethics approval

Peripheral blood mononuclear cells (PBMC) were isolated from buffy coats obtained from blood donations. Blood samples were drawn with the informed consent of the human subjects. The study was approved by the Ethical Committee of the University of Wuppertal.

## Source of biological material

PBMC were isolated from buffy coats obtained from blood donations at the University Hospital Düsseldorf and Deutsches Rote Kreuz-blood donation service West (Hagen).

## Abbreviations

ARA: arachidonic acid
COX: cyclooxygenase
DHA: docosahexaenoic acid
diHEPE: dihydroxy eicosapentaenoic acid
diHDHA: dihydroxy docosahexaenoic acid
diHDPE: dihydroxy docosapentaenoic acid
diHETE: dihydroxy eicosatetraenoic acid
EP: prostaglandin E_2_ receptor
EPA: eicosapentaenoic acid
EpDPE: epoxy docosapentaenoic acid
EpETE: epoxy eicosatrienoic acid
ESI: electrospray ionization
FA: fatty acid
GM-CSF: granulocyte-macrophage colony-stimulating factor
HDHA: hydroxy docosahexaenoic acid
HEPE: hydroxy eicosapentaenoic acid
HETE: hydroxy eicosatetraenoic acid
HUFA: highly unsaturated fatty acid
IS: internal standard
LOX: lipoxygenase
M-CSF: macrophage colony-stimulating factor
MS: mass spectrometry
MUFA: monounsaturated fatty acid
PBMC: peripheral blood mononuclear cell
PG: prostaglandin
P/S: penicillin/streptomycin
PUFA: polyunsaturated fatty acid
Rv: resolvin
SFA: saturated fatty acid
Tx: thromboxane

## Supplementary material

**Figure S1:**
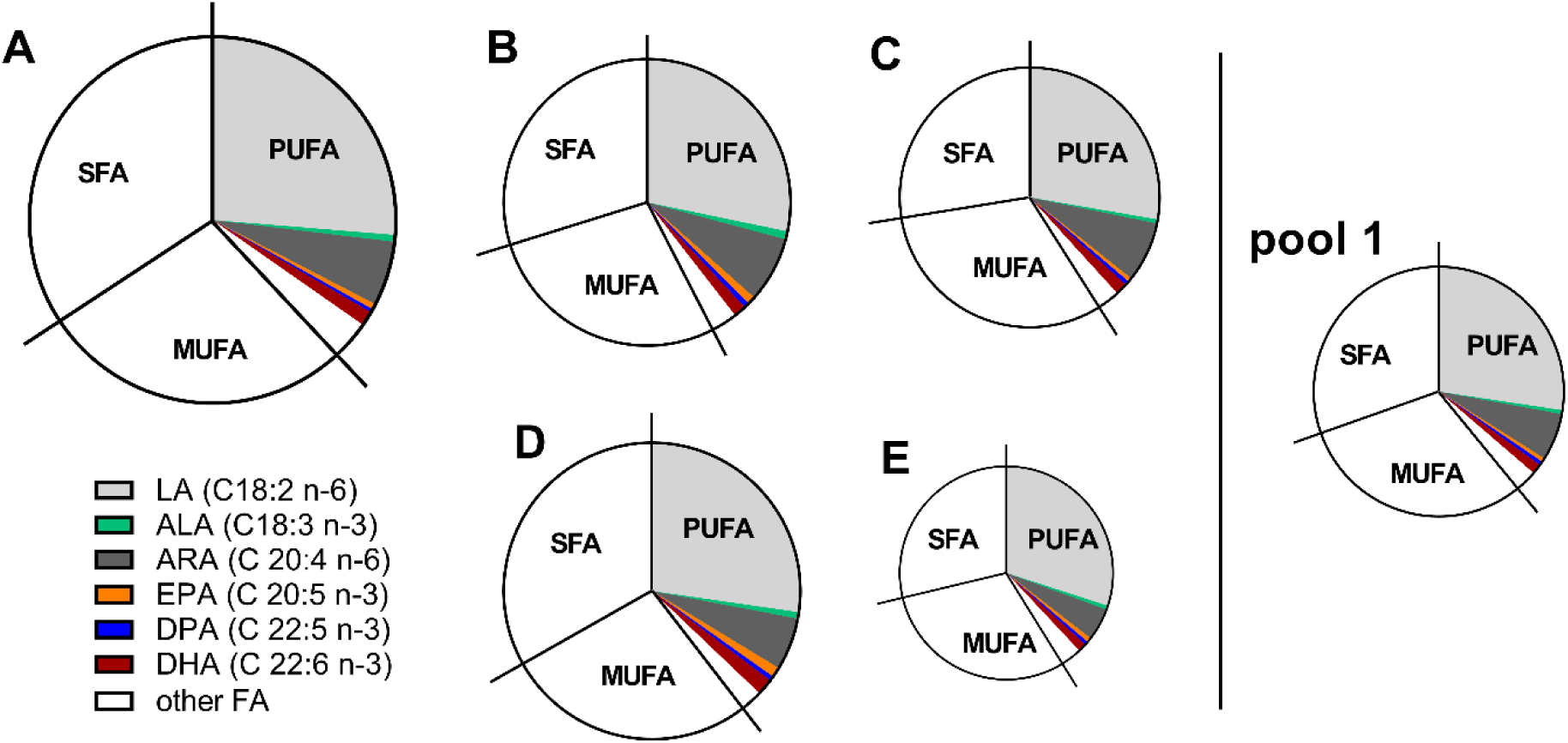
Relative fatty acid profiles of the tested human citrate plasmas. Non-fasting plasma of five subjects from a local blood donation center was analyzed for total FA concentrations by means of LC-MS/MS. Plasmas C and E were pooled resulting in plasma pool 1. Sizes of the circles indicate relative total fatty acid concentrations (plasma A, 11.0 ± 0.7 mM; plasma B, 8.7 ± 0.7 mM; plasma C, 7.8 ± 0.5 mM; plasma D, 9.0 ± 0.7 mM; plasma E, 6.4 ± 0.4 mM; pool 1, 7.5 ± 0.7 mM). SFA, saturated fatty acid; MUFA, monounsaturated fatty acid; PUFA, polyunsaturated fatty acid.

**Table S1:**
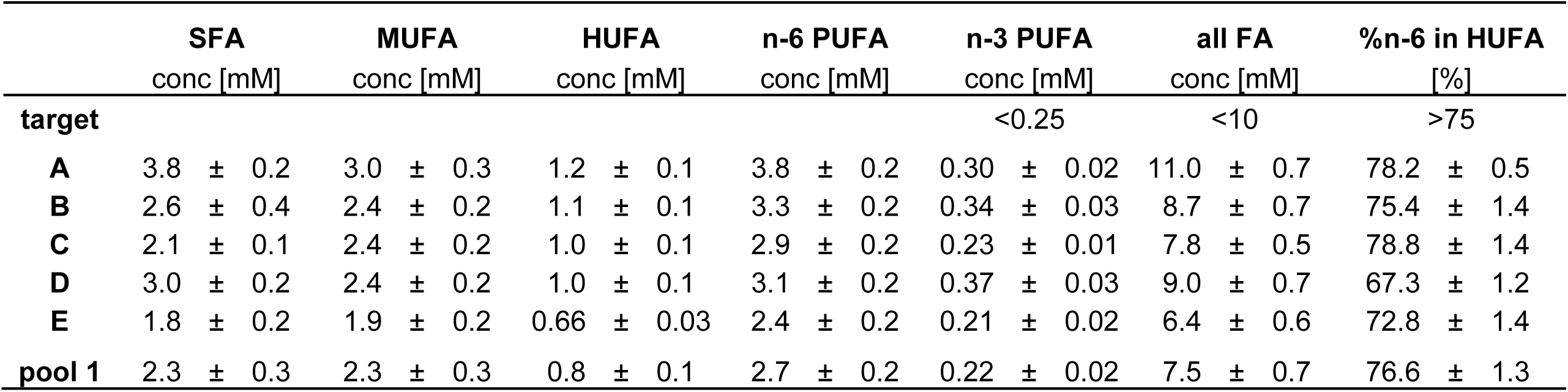
Fatty acid pattern of the tested human citrate plasmas. Non-fasting plasma of five subjects (A-E) from a local blood donation center and plasma pool 1 (pooled plasma of C and E) was analyzed for total FA concentrations by means of LC-MS/MS. Results are shown as mean ± SD, *n* = 3. %n-6 in HUFA was calculated from fatty acid concentrations of C20:3 n-6, C20:4 n-6, C22:4 n-6, C22:5 n-6, C20:3 n-9, C20:5 n-3, C22:5 n-3, C22:6 n-3.

|  | **SFA**<br>conc [mM] | **MUFA**<br>conc [mM] | **HUFA**<br>conc [mM] | **n-6 PUFA**<br>conc [mM] | **n-3 PUFA**<br>conc [mM] | **all FA**<br>conc [mM] | **%n-6 in HUFA**<br>[%] |
| --- | --- | --- | --- | --- | --- | --- | --- |
| **target** |  |  |  |  | <0.25 | <10 | >75 |
| **A** | 3.8 ± 0.2 | 3.0 ± 0.3 | 1.2 ± 0.1 | 3.8 ± 0.2 | 0.30 ± 0.02 | 11.0 ± 0.7 | 78.2 ± 0.5 |
| **B** | 2.6 ± 0.4 | 2.4 ± 0.2 | 1.1 ± 0.1 | 3.3 ± 0.2 | 0.34 ± 0.03 | 8.7 ± 0.7 | 75.4 ± 1.4 |
| **C** | 2.1 ± 0.1 | 2.4 ± 0.2 | 1.0 ± 0.1 | 2.9 ± 0.2 | 0.23 ± 0.01 | 7.8 ± 0.5 | 78.8 ± 1.4 |
| **D** | 3.0 ± 0.2 | 2.4 ± 0.2 | 1.0 ± 0.1 | 3.1 ± 0.2 | 0.37 ± 0.03 | 9.0 ± 0.7 | 67.3 ± 1.2 |
| **E** | 1.8 ± 0.2 | 1.9 ± 0.2 | 0.66 ± 0.03 | 2.4 ± 0.2 | 0.21 ± 0.02 | 6.4 ± 0.6 | 72.8 ± 1.4 |
| **pool 1** | 2.3 ± 0.3 | 2.3 ± 0.3 | 0.8 ± 0.1 | 2.7 ± 0.2 | 0.22 ± 0.02 | 7.5 ± 0.7 | 76.6 ± 1.3 |

**Table S2:** Oxidation status of the tested human citrate plasmas. Non-fasting plasma of five subjects (A-E) from a local blood donation center and plasma pool 1 (pooled plasma of C and E) was analyzed for total oxylipin concentrations by means of LC-MS/MS. Shown are concentrations of representative oxylipins formed by autoxidation and/or enzymatic activity as mean ± SD, *n* = 3.

|  | target | plasma A | plasma B | plasma C | plasma D | plasma E | pool 1 |
| --- | --- | --- | --- | --- | --- | --- | --- |
|  | [nM] | conc [nM] | conc [nM] | conc [nM] | conc [nM] | conc [nM] | conc [nM] |
| **5-HETE** | <100 | 17.6 ± 0.6 | 17.3 ± 0.9 | 13.6 ± 0.8 | 15 ± 1 | 9 ± 0.5 | 12 ± 1 |
| **12-HETE** | <200 | 17 ± 1 | 22 ± 1 | 14.5 ± 0.6 | 13.3 ± 0.1 | 13 ± 1 | 15 ± 1 |
| **15-HETE** | <200 | 21 ± 2 | 24 ± 2 | 16.8 ± 0.6 | 15.2 ± 0.6 | 9.5 ± 0.9 | 14.3 ± 0.9 |
| **5-F2t-IsoP** | <1 | 0.43 ± 0.04 | 0.52 ± 0.07 | 0.21 ± 0.05 | 0.51 ± 0.12 | 0.31 ± 0.06 | 0.35 ± 0.03 |
| **9-HODE** | <500 | 137 ± 9 | 137 ± 7 | 197 ± 13 | 214 ± 5 | 79 ± 7 | 137 ± 10 |

**Table S3:** Fatty acid status of erythrocytes from the human subjects who donated the monocytes used for n-3 PUFA supplementation experiments. Erythrocytes were analyzed for total fatty acid concentrations by means of LC-MS/MS and %n-6 in HUFA was calculated from fatty acid concentrations of C20:3 n-6, C20:4 n-6, C22:4 n-6, C22:5 n-6, C20:3 n-9, C20:5 n-3, C22:5 n-3, C22:6 n-3. All donors were healthy, voluntary blood donors from local blood donation centers. Results are shown as mean ± SD, *n* = 3.

| experi-<br>ment | subject | %n-6 in HUFA | LA (C18:2 n-6) | ARA (C20:4 n-6) | EPA (C20:5 n-3) | DHA (C22:6 n-3) | DPA (C22:5 n-3) |
| --- | --- | --- | --- | --- | --- | --- | --- |
| | | [%] | conc [μM] | conc [μM] | conc [μM] | conc [μM] | conc [μM] |
| 1 | 1 | 76.8 ± 0.6 | 626 ± 93 | 699 ± 48 | 36 ± 3 | 148 ± 16 | 112 ± 11 |
| | 2 | 74.6 ± 0.6 | 914 ± 184 | 745 ± 93 | 31 ± 5 | 188 ± 31 | 116 ± 17 |
| | 3 | 74.7 ± 0.4 | 538 ± 20 | 842 ± 56 | 37 ± 2 | 207 ± 9 | 135 ± 5 |
| | 4 | 71.1 ± 0.3 | 616 ± 40 | 683 ± 47 | 35 ± 3 | 221 ± 12 | 113 ± 6 |
| 2 | 5 | 75.4 ± 1.0 | 764 ± 29 | 674 ± 23 | 26 ± 3 | 222 ± 24 | 86 ± 6 |
| | 6 | 70.0 ± 0.6 | 844 ± 41 | 585 ± 45 | 50 ± 3 | 220 ± 9 | 106 ± 5 |
| | 7 | 76.2 ± 0.3 | 751 ± 4 | 709 ± 17 | 31 ± 1 | 187 ± 4 | 105 ± 5 |
| | 8 | 75.9 ± 0.6 | 1322 ± 117 | 753 ± 86 | 44 ± 4 | 166 ± 15 | 136 ± 7 |

**Figure S2:**
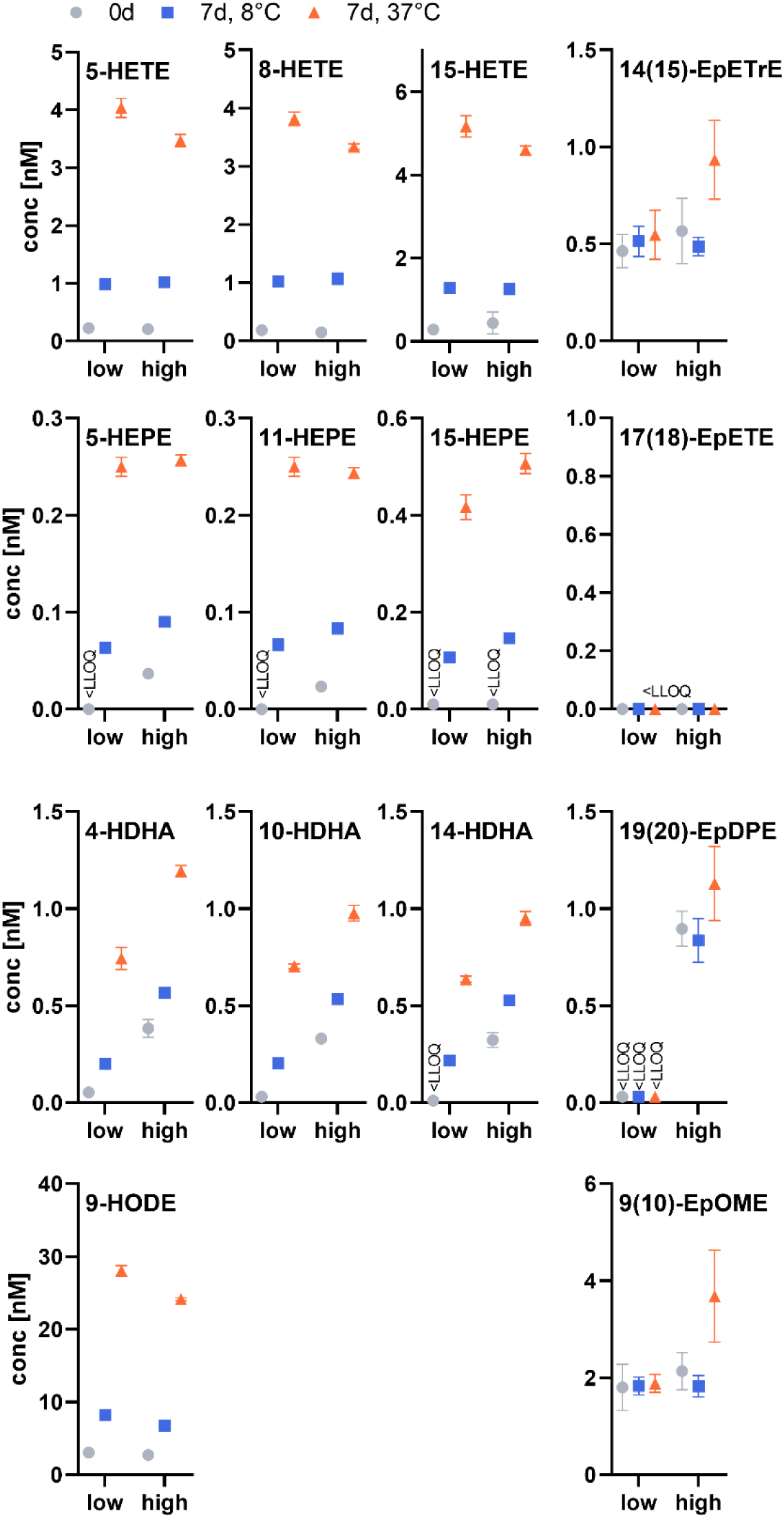
Oxylipin concentrations of the used cell culture media at baseline and after storage for 7 days. Media were prepared with 5% (*v*/*v*) plasma pool 1 (non-supplemented medium, low). For the supplemented medium (high), additionally 10.3 µM DHA (>99%) and 5.5 µM EPA (>99%) were added. Concentrations of selected, most abundant oxylipins derived from ARA, EPA, DHA and LA were analyzed at beginning and after 7 days of incubation at 8 °C or 37 °C by means of LC-MS/MS. Results of one representative experiment are shown as mean ± SD, *n* = 3.

**Figure S3:**
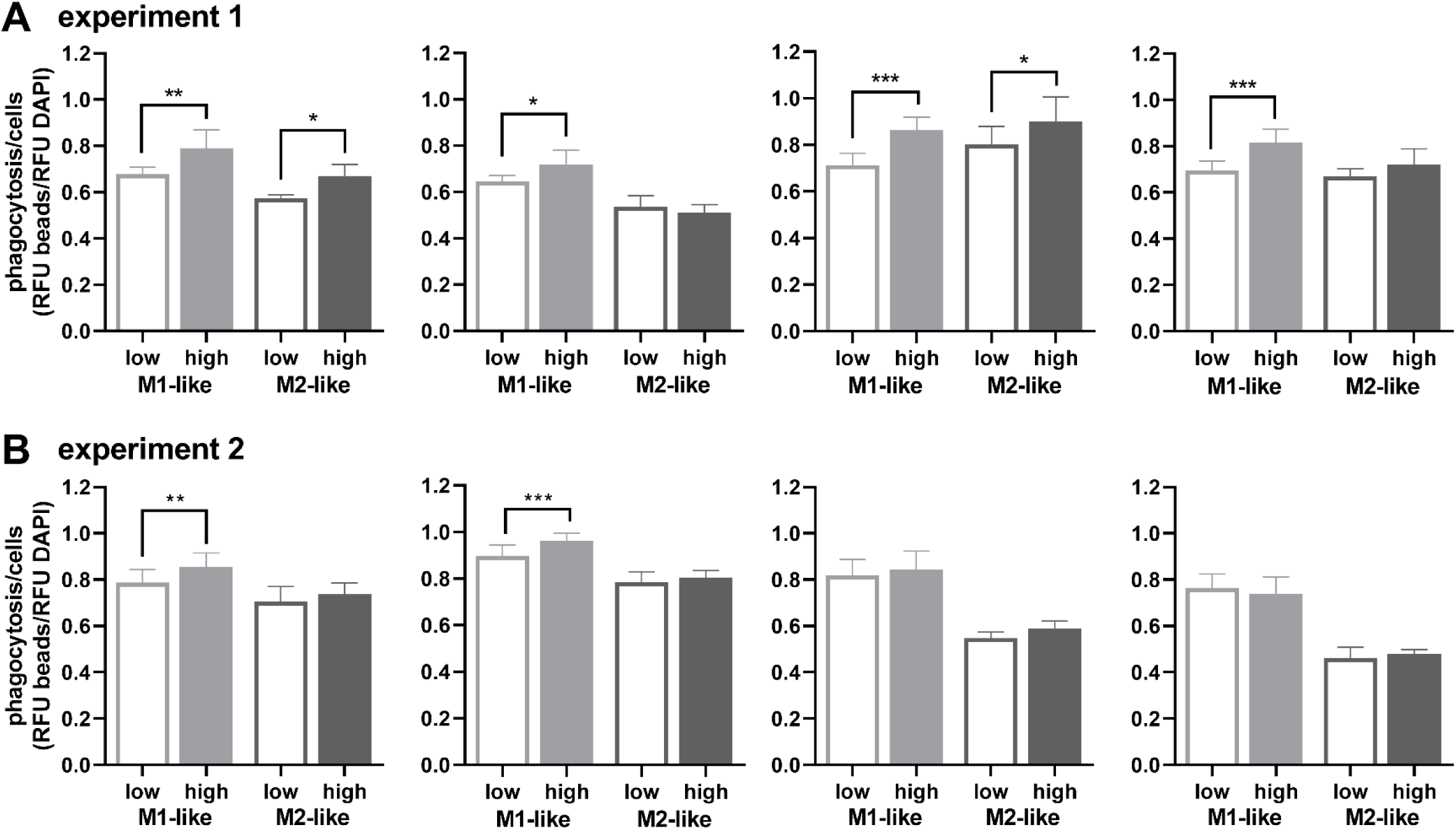
Impact of n-3 PUFA supplementation on phagocytosis in the two independent supplementation experiments (A, B). After isolation of monocytes, cells were differentiated into macrophages and supplemented using medium with 5% (*v*/*v*) plasma pool 1 (low) or plasma pool 1 and 10.3 µM DHA (>99%) and 5.5 µM EPA (>99%) (high) for 2 days. Cells were transferred into 96 well plates (50,000 cells per well) for phagocytosis assay. Shown is phagocytosis as ratio of bead fluorescence to DAPI fluorescence. Results of four independently performed experiments are shown as mean ± SD for *n* = 5-8 (A) or *n* = 14-15 (B) replicates from a pool of 4 subjects. Statistical analysis was performed by 1-way ANOVA followed by Sidak’s multiple comparison test. Differences from vehicle control were considered significant at *p* values ≤0.5 (*), ≤0.01 (**) or ≤0.001 (***).

**Table S4:** Concentrations of oxylipin standards (stock solutions) used for supplementation of the medium analyzed by means of LC-MS/MS.

| mix | analyte | concentration |  |
| --- | --- | --- | --- |
|  |  | nominal<br>[mM] | measured<br>[mM] |
| I | 4-HDHA | 0.29 | 0.25 |
|  | 7-HDHA | 0.29 | 0.32 |
|  | 5-HEPE | 0.31 | 0.18 |
| II | 14(S)-HDHA | 0.29 | 0.21 |
|  | 17(S)-HDHA | 0.29 | 0.22 |
|  | 12(S)-HEPE | 0.31 | 0.41 |
|  | 15(S)-HEPE | 0.31 | 0.42 |
| III | 19(20)-EpDPE | 0.29 | 0.28 |
|  | 17(18)-EpETE | 0.31 | 0.43 |

**Figure S4:**
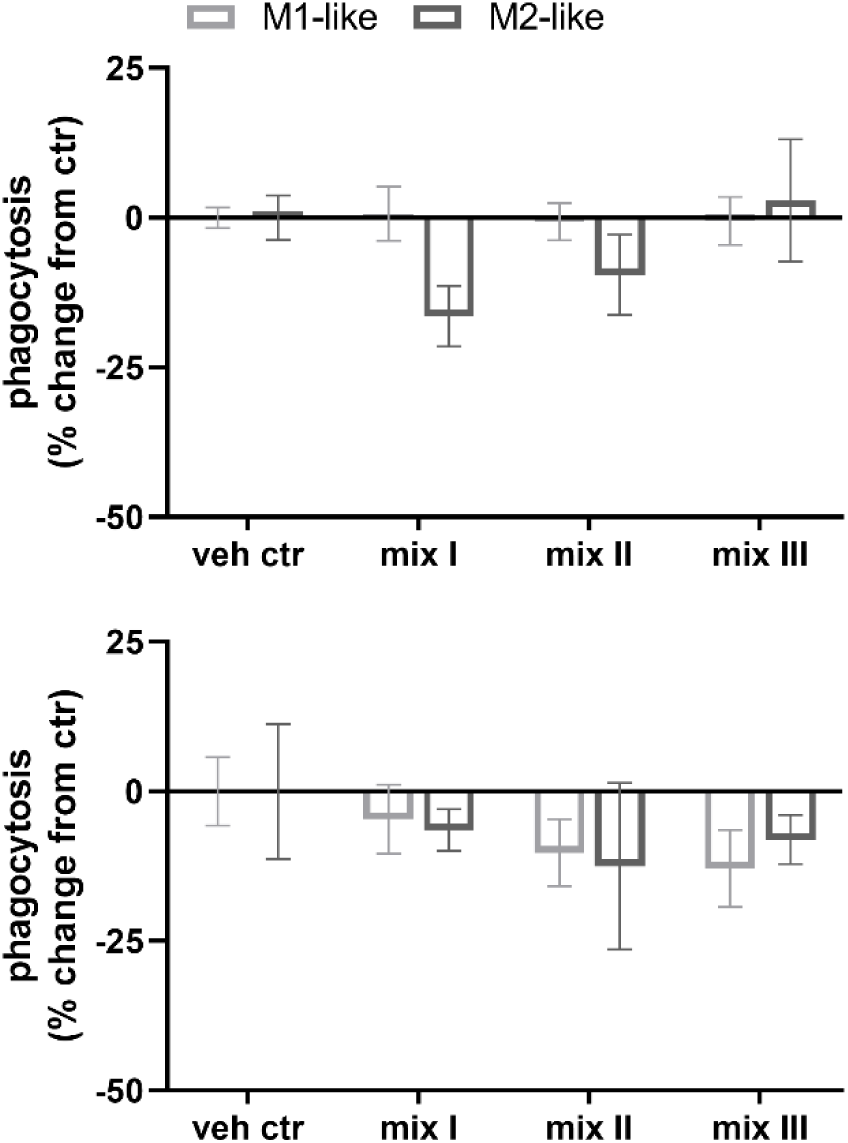
Impact of mixtures of oxylipins on phagocytosis. Non-supplemented macrophages were transferred into 96 well plates (50,000 cells per well) and incubated with 300 nM oxylipins (mix I: 5-HEPE, 4-HDHA, 7-HDHA; mix II: 12(*S*)-, 15(*S*)-HEPE, 14(*S*)-,17(*S*)-HDHA; mix III: 17(18)-EpETE, 19(20)-EpDPE, Table S4) or 0.1% DMSO (veh ctr) for 1 h. Shown are the results of two independently performed phagocytosis experiments as % change from vehicle control as mean ± SD for *n* = 4-5 replicates from a pool of 4 subjects.

**Table S5:** Concentrations of added oxylipins in the media after phagocytosis assay. Macrophages were preincubated with a mix of oxylipins (300 nM) for 1 h. Phagocytosis was started by addition of fluorescent beads and the mix of oxylipins was renewed. After 2 h the medium was collected and analyzed for non-esterified oxylipins by means of LC-MS/MS. Results of one representative experiment are shown as mean ± SD, *n* = 3.

| mix | oxylipin | M1-like |  |  | M2-like |  |  |
| --- | --- | --- | --- | --- | --- | --- | --- |
|  |  | conc |  |  | conc |  |  |
|  |  | [nM] |  |  | [nM] |  |  |
| I | 5-HEPE | 246 | ± | 10 | 173 | ± | 7 |
| | 4-HDHA | 162 | ± | 6 | 124 | ± | 9 |
| | 7-HDHA | 189 | ± | 5 | 161 | ± | 8 |
| II | 12-HEPE | 125 | ± | 1 | 105 | ± | 4 |
| | 15-HEPE | 143 | ± | 2 | 115 | ± | 3 |
| | 14-HDHA | 128 | ± | 4 | 110 | ± | 3 |
| | 17-HDHA | 204 | ± | 1 | 158 | ± | 7 |
| III | 17(18)-EpETE | 232 | ± | 8 | 159 | ± | 7 |
| | 17,18-DiHETE | 18 | ± | 1 | 26 | ± | 2 |
| | 19(20)-EpDPE | 294 | ± | 13 | 238 | ± | 5 |
| | 19,20-DiHDPE | 14 | ± | 1 | 21 | ± | 1 |

**Table S6:** Concentrations of added oxylipins in the media after phagocytosis assay. Macrophages were preincubated with 100 nM PGE_2_ (control) or additionally a mix of oxylipins (300 nM) for 1 h. Phagocytosis was started by addition of fluorescent beads and the mix of oxylipins was renewed. After 2 h the medium was collected and analyzed for non-esterified oxylipins by means of LC-MS/MS. Results of one representative experiment are shown as mean ± SD, *n* = 3.

| mix | oxylipin | M1-like | M2-like |
| --- | --- | --- | --- |
|  |  | conc<br>[nM] | conc<br>[nM] |
| control | PGE<sub>2</sub> | 90 ± 2 | 96 ± 1 |
| I | 5-HEPE | 12.5 ± 0.4 | 21 ± 1 |
| | 4-HDHA | 10 ± 1 | 14 ± 1 |
| | 7-HDHA | 12 ± 1 | 17 ± 1 |
| | PGE<sub>2</sub> | 86 ± 4 | 88 ± 2 |
| II | 12-HEPE | 7.9 ± 0.4 | 10 ± 1 |
| | 15-HEPE | 8.9 ± 0.4 | 12 ± 1 |
| | 14-HDHA | 8.2 ± 0.4 | 10 ± 1 |
| | 17-HDHA | 14 ± 2 | 21 ± 2 |
| | PGE<sub>2</sub> | 84 ± 2 | 91 ± 2 |
| III | 17(18)-EpETE | 17 ± 1 | 22 ± 2 |
| | 17,18-DiHETE | 4.1 ± 0.1 | 2.8 ± 0.2 |
| | 19(20)-EpDPE | 25 ± 1 | 29 ± 1 |
| | 19,20-DiHDPE | 3.0 ± 0.1 | 1.7 ± 0.2 |
| | PGE<sub>2</sub> | 87 ± 5 | 92 ± 9 |

**Figure S5:**
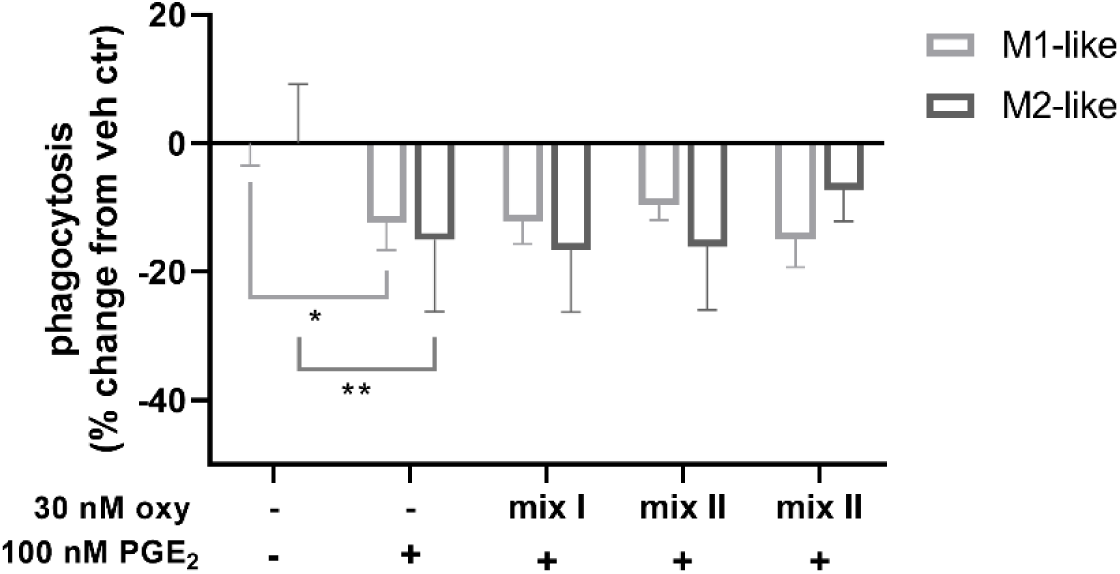
Impact of n-3 PUFA derived oxylipins on the inhibitory effect of PGE_2_ on phagocytosis. Primary human macrophages were preincubated with 100 nM PGE_2_ and 30 nM oxylipins (mix I: 5-HEPE, 4-HDHA, 7-HDHA; mix II: 12(*S*)-, 15(*S*)-HEPE, 14(*S*)-, 17(*S*)-HDHA; mix III: 17(18)-EpETE, 19(20)-EpDPE, Table S4) or 0.1% DMSO (veh ctr) for 1 h, followed by phagocytosis for 2 h. Shown is phagocytosis as % change from vehicle control as mean ± SD for *n* = 4-5 replicates from a pool of 4 subjects. Statistical analysis was performed by 1-way ANOVA followed by Sidak’s multiple comparison test. Differences from vehicle control were considered significant at *p* values ≤0.5 (*), ≤0.01 (**).

